# Cell-penetrating peptides stimulate protein transport on the Twin-arginine translocation pathway

**DOI:** 10.1101/2023.07.08.548235

**Authors:** Robert McNeilage, Iniyan Ganesan, Johnathan Keilman, Steven M. Theg

## Abstract

The Tat pathway is essential for photosynthetic protein transport across plant thylakoid membranes and is also ubiquitous throughout prokaryotes and archaea. The Tat pathway is unique amongst protein translocation pathways as it specializes in transporting folded proteins driven by a proton motive force. Mechanistic details of the actual translocation step (s) of the pathway remain elusive. Here, we show that membrane thinning stimulates Tat transport and, conversely, membrane strengthening abolishes Tat transport. We draw parallels from the Tat transport mechanism to that of cell penetrating peptides and propose that the Tat pore could be toroidal in shape and lined by lipids, as in those formed by cell penetrating peptides.

**Significance Statement:** Protein translocation across membranes is a significant cellular activity in both prokaryotes and eukaryotes. The Tat pathway for protein translocation operates in bacteria, archaea, chloroplasts, and plant mitochondria. Its mechanism of action has been difficult to decipher, but recent evidence suggests it does not use a conical proteinaceous transport channel. Instead, it has been suggested to translocate proteins through lipid-lined toroidal pores set up by membrane thinning. This work supports that hypothesis by showing that membrane-thinning cell-penetrating peptides stimulate the Tat pathway in both chloroplasts and bacterial plasma membranes, and that membrane stabilization blocks the pathway. We believe this is the most direct evidence to date of the toroidal pore mechanism operating in the Tat pathway.

## Introduction

The Twin-Arginine Translocation (Tat) pathway transports folded proteins across the thylakoid membrane of chloroplasts, the inner membrane in plant mitochondria, and the periplasmic membranes of bacteria and archaea utilizing a proton motive force (pmf) as its sole energy source (Hamsanathan & Musser, 2018, New et al., 2018, Palmer & Stansfeld, 2020). Because both photosynthetic proteins and bacterial virulence factors(De Buck et al., 2008) pass through the Tat pathway, it is an important process to understand from multiple human health perspectives, both in nutrition and disease. In most instances, Tat transport is mediated by three proteins: Tha4, Hcf106, and cpTatC in thylakoids; TatA, TatB, and TatC in bacteria. Tha4 and Hcf106 are structurally similar, being composed of an N-terminal transmembrane helix (TMH), followed by a hinge region, an amphipathic helix (APH) that lies along the surface of the membrane, and an unstructured C-terminal domain. The Hcf106 APH and C-terminal domains are longer than their Tha4 counterparts(Porcelli et al., 2002). Interestingly, the NMR structures of TatA and TatB show that the TMHs are shorter than the hydrophobic portion of the lipid bilayer(Rodriguez et al., 2013a, Zhang et al., 2014), which creates a hydrophobic mismatch. TatC has six transmembrane domains with a topology where both termini face the stroma(Gouffi et al., 2002, Rollauer et al., 2012). Two of the TatC TMHs are also short, again leading to hydrophobic mismatch(Ramasamy et al., 2013, Rollauer et al., 2012). This hydrophobic mismatch has recently been shown to be important for the efficient operation of the Tat pathway(Hao et al., 2022a, Mehner-Breitfeld et al., 2022, Stockwald et al., 2022). Translocation of proteins along this pathway requires the precursor protein to possess an N-terminal signal peptide that enables the precursor to be recognized and bound by the receptor complex. The signal peptide contains the essential and eponymous RR (twin arginine) motif(Stanley et al., 2000). Oligomers of Hcf106 and TatC in a 1:1 ratio compose the receptor complex for binding of the precursor signal sequence(Bolhuis et al., 2001) in an energy independent manner(Mori & Cline, 2002). After precursor binding and in the presence of a pmf, the receptor complex recruits multimers of Tha4 and translocation occurs(Mori & Cline, 2002). After the translocation event, the active translocon dissociates into the Hcf106-cpTatC receptor complex and oligomers of Tha4(Mori & Cline, 2002). Sometime during or after the translocation event, the signal peptide is cleaved. The detailed mechanism of the actual translocation event following Tha4 recruitment is not yet understood.

Currently, there are three competing models proposed to accomplish protein transport on the Tat pathway: the protein iris model in which Tha4 joins the translocon in varying stoichiometry to form a tightly sealing pore around the incoming protein, the co-enzyme model, in which TatA is thought to act as a co-enzyme that accumulates at the translocation site to activate a translocon built from a complex consisting of TatB and TatC (Hauer et al., 2017, Hauer et al., 2013), and the membrane defect model, in which the membrane is proposed to form toroidal pores in response to bilayer destabilizing forces and through which proteins traverse the membranes. A recent variation of the latter model posits that the transmembrane helix (TMH) of TatA flips to lie along the trans-side of the membrane surface, contributing to the bilayer instability (Stockwald et al., 2022).

The protein iris model faces several structural challenges: the pore would be either lined with the hydrophobic TMHs, or the APHs of Tha4 would have to fold back in a hairpin onto the TMDs. The hairpin fold is not favored by experimental evidence(Alcock et al., 2017, Aldridge et al., 2012, Koch et al., 2012). Despite the Tat pathway translocating precursors of different shapes and sizes, the membranes of the thylakoids remain tightly sealed during translocation and do not allow for leakage of ions, which seems unlikely with the proteinaceous pores just described(Asher & Theg, 2021, Teter & Theg, 1998). These findings, coupled with the transient assembly of the active translocon, have led to the suggestion that the Tat pathway does not utilize a proteinaceous pore to achieve translocation(Asher & Theg, 2021, Bruser & Sanders, 2003, Hao et al., 2022a). It has been proposed that instead, under the influence of the Tat machinery subunits, the transported substrate, and membrane energization the membrane is locally thinned to an extent that defects in the bilayer develop that allow a precursor to pass through the membrane itself via lipid lined toroidal pores (reviewed in(Patel et al., 2014)). A key component in this model is the thylakoid membrane thinning that was observed almost 50 years ago to occur in the light (Murakami & Packer, 1970a), and which has recently been repeated in two different laboratories (Johnson et al., 2011a, Johnson et al., 2011b, Kirchhoff et al., 2011, Li et al., 2020). We believe this provides a compelling mechanism for coupling between the pmf and cpTat transport as the former causes membrane thinning and the latter depends on it.

Cell Penetrating Peptides (CPPs) are short amphipathic peptides that interact with and can induce toroidal pore formation in membranes (Huang & Li, 2023, Leveritt Iii et al., 2015). Many CPPs adopt an α-helical configuration when interacting with membranes. The α-helices are usually amphipathic, with positively charged residues, often arginine, on one side of the helix and hydrophobic residues on the other side. The hydrophobic residues interface with the fatty acid side chains, while the positively charged residues interact with the polar or negatively charged lipid head groups of membranes. CPPs have generated considerable interest for their ability to spontaneously cross membranes by causing membrane thinning and the formation of toroidal pores(Herce & Garcia, 2007, Huang & Li, 2023, Madani et al., 2011). The ability to thin membranes to the point of pore formation by CPPs suggests a possible mechanism for Tat protein translocation, where the potential membrane thinning effect of the pmf combined with destabilizing effects of the Tat component subunits, i.e., hydrophobic mismatch, might lead to local and transient toroidal pore formation.

In this study, we used CPPs to initiate the thinning of thylakoid and *E. coli* membranes(Chen et al., 2003, Mecke et al., 2005). We explored the idea that the CPPs can provide some of the energy normally generated by the pmf to the Tat system by pre-thinning the membrane. We demonstrate that CPPs specifically stimulate Tat transport and that the opposite effect of strengthening the membrane abolishes transport. We believe that these experiments establish the central role of membrane thinning in the Tat mechanism.

## Results

### Cell Penetrating Peptides stimulate Tat protein translocation

The mechanism through which CPPs cross membranes is proposed to involve actively thinning the membrane to the point that the bilayer breaks down with the formation of toroidal pores(Chen et al., 2003, Herce & Garcia, 2007). If the Tat pathway operates under similar principles, that is, through membrane thinning, then we reasoned that CPPs should stimulate the transport of proteins on the Tat pathway by contributing to the membrane thinning process. To test this hypothesis, different CPPs were added to thylakoid Tat pathway transport reactions. We chose four peptides from unrelated sources with widely differing sequences and related only in their ability to spontaneously penetrate membranes. Fig. 1 shows that each peptide tested stimulated the transport of iOE17, a model Tat substrate in thylakoids(Alder & Theg, 2003).

**Figure 1.**
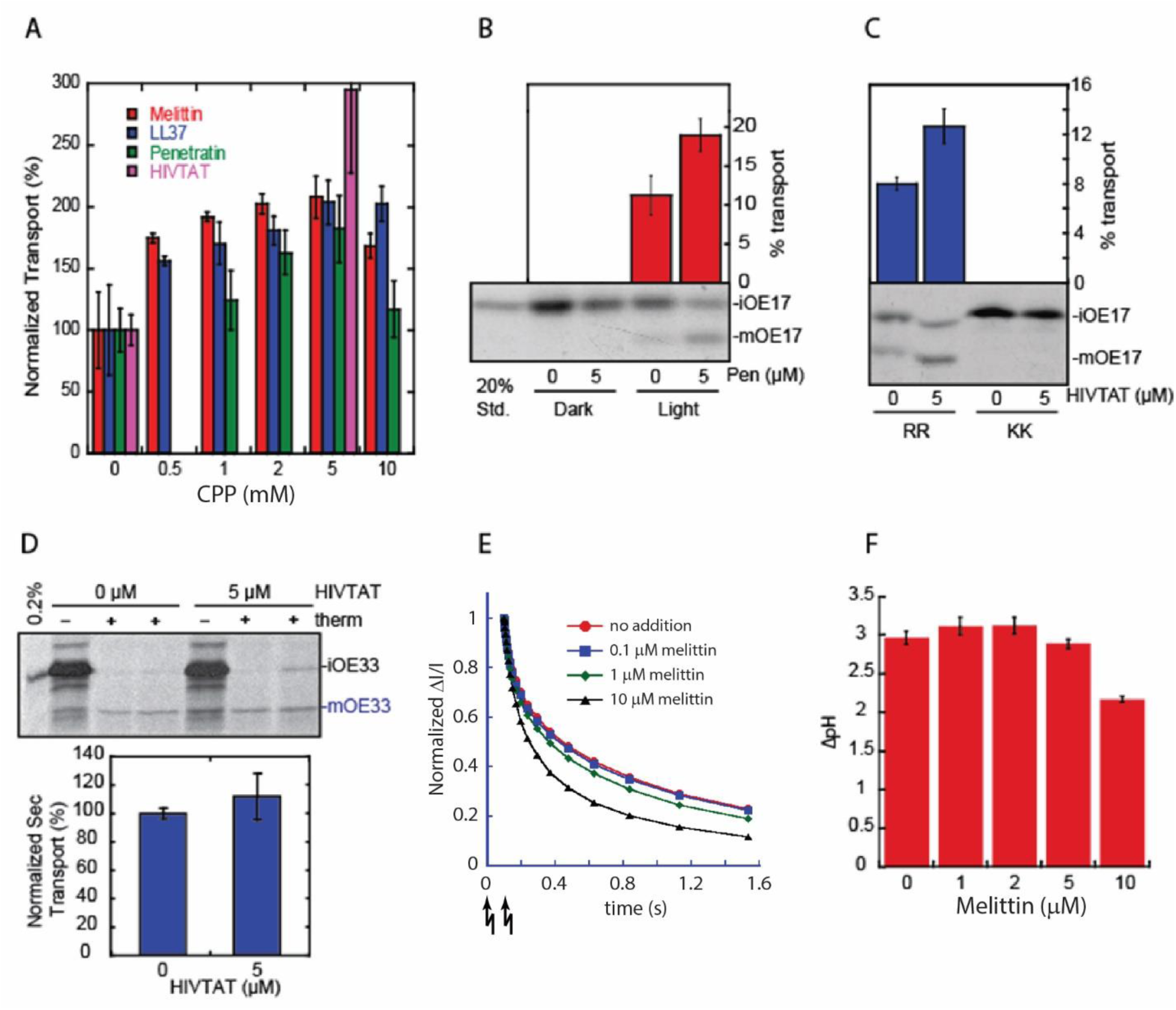
CPPs specifically stimulate transport on the Tat pathway. In A-D, thylakoids were incubated with a radiolabeled substrate and CPPs at the indicated concentrations. A) Thylakoids were treated with CPPs during a transport reaction of iOE17. Quantification of the mature protein is shown. B) Thylakoids were incubated in the dark for 30 min (D) or light (L) with iOE17 and with or without 5 μM penetratin. C) Thylakoids were treated with given HIVTat concentrations and mixed with the transport incompetent precursor, iOE17KK. D) Thylakoids were treated with HIVTat as indicated and assayed for transport of iOE33 through the Sec pathway. Post-transport thylakoids were treated with thermolysin (therm) where indicated. E) ECS measurements were made on 80 μg thylakoids with the indicated concentrations of melittin. Two 9 ms-duration flashes were delivered 100 ms apart (arrows at the bottom of the panel); the ECS signal was normalized to the first data point after the second flash. F) ΔpH measurements were made by observing the quenching of NED fluorescence induced by application of actinic light with indicated concentrations of melittin.

Because CPPs are known to form pores in membranes(Herce & Garcia, 2007), we tested the possibility of passive diffusion of precursors into the thylakoids through a CPP induced pore. Additionally, CPPs are the subject of investigation for delivering drugs and proteins across membranes(Ramsey & Flynn, 2015), therefore, we address the possibility of precursors hitchhiking on CPPs simultaneously with the previous concern. To these ends, we asked whether CPPs could facilitate transport either using a transport-incompetent precursor or in the absence of an energy source. If CPPs promote spontaneous diffusion of precursors across thylakoids membranes, then they should cause translocation in the absence of a pmf. Thylakoids, CPPs, and iOE17 were incubated together in the dark at ambient temperature for 30min. CPPs did not allow for translocation without the light-generated pmf (Fig. 1B). In the presence of a light source, translocation occurred and was stimulated by CPPs. We then asked if the pmf and CPPs would be able to cause the translocation of a transport-incompetent Tat substrate. We used a mutated iOE17 with RR to KK substitutions in the signal sequence (iOE17KK), which renders the precursor transport-incompetent(Ize et al., 2002). iOE17KK was not transported with or without CPPs present (Fig 1C). These results preclude the possibility that either precursors or the thylakoid processing protease can either diffuse through pores created by CPPs or be carried by them across the thylakoid membrane in a Tat-independent or energy-independent manner.

In these two controls, the precursor was found with reisolated thylakoids. This indicates that being present at the membrane is not sufficient for CPPs to facilitate passive transport of precursors. These results also suggest that if precursors travel through any pore in the membrane, it was induced and controlled by the Tat machinery, which would have been inactive in these controls.

To determine the specificity of CPP stimulation of the Tat pathway, we monitored the effect of CPPs on protein translocation across thylakoids via the Sec pathway, which occurs through a proteinaceous pore (Li et al., 2016). Transport of the model thylakoid Sec substrate, iOE33, was not significantly altered by HIVTat (Fig. 1D). This clearly establishes that CPPs do not universally stimulate protein translocation.

Figure 1a shows that stimulation of the Tat pathway drops at higher concentrations of melittin and penetratin. We reasoned that these CPPs caused enough ion leakage to overwhelm the ability of the thylakoids to maintain the pmf, leading to the decline in transport stimulation. To investigate this possibility, we monitored the effect of higher concentrations of melittin on both the electrical and chemical components of the pmf. Melittin at 10 μM caused a faster dissipation of the light-induced Δψ across the thylakoid membrane compared with controls (Fig. 1E). Similarly, melittin at 10 µM resulted in a reduced steady-state trans-membrane ΔpH (Fig. 1F). Both of these observations are consistent with CPPs mediating proton and other ion leakage through pores induced at these higher concentrations. Taken together, these results indicate that high concentrations of CPPs compromise the integrity of the thylakoid membranes, likely by inducing transient pores(Herce & Garcia, 2007). This is not surprising because inducing pores in and causing the rupturing of membranes is a well-documented effect of some CPPs(Herce & Garcia, 2007, Yang et al.), leading them also to be described as antimicrobial peptides (Huang & Li, 2023).

### Effects of CPPs on the energetics of Tat protein transport

Next, we further evaluated the mechanism of CPP-mediated Tat transport stimulation. We reasoned that the light- and pH-induced membrane thinning of thylakoids observed in the literature(Kirchhoff et al., 2011, Li et al., 2020, Murakami & Packer, 1970b) could provide a mechanism through which the pmf could be coupled to protein translocation on the Tat pathway if the latter depended on membrane thinning. This would also provide an explanation of the stimulatory effect of CPPs, since they are proposed to act via membrane thinning(Herce & Garcia, 2007, Madani et al., 2011). Specifically, if both CPPs and the pmf contribute to Tat protein transport through their effects on membrane thickness, then we would expect that CPPs would lower the threshold energy needed to promote protein transport by providing some thinning before application of the ΔpH. To test this idea, we measured translocation as a function of ΔpH to elucidate the threshold ΔpH at which translocation begins(Rollauer et al., 2012).

In chemiosmotic systems, plotting the work performed by the pmf as a function of the driving force results in a biphasic plot. Initially, there is not enough energy in the gradient to perform the work queried. At some point a threshold is reached, above which the output rises linearly with the driving force. This thermodynamic threshold represents the minimum energy required for the work queried, in this case protein transport (Alder & Theg, 2003). Figure 2 shows that the stimulatory effect of both melittin and penetratin is caused by their lowering of the ΔpH threshold for Tat protein transport. This is consistent with the idea that the CPPs predispose the membranes for protein transport by mediating part of the essential thinning that is then finished by the pmf.

**Figure 2.**
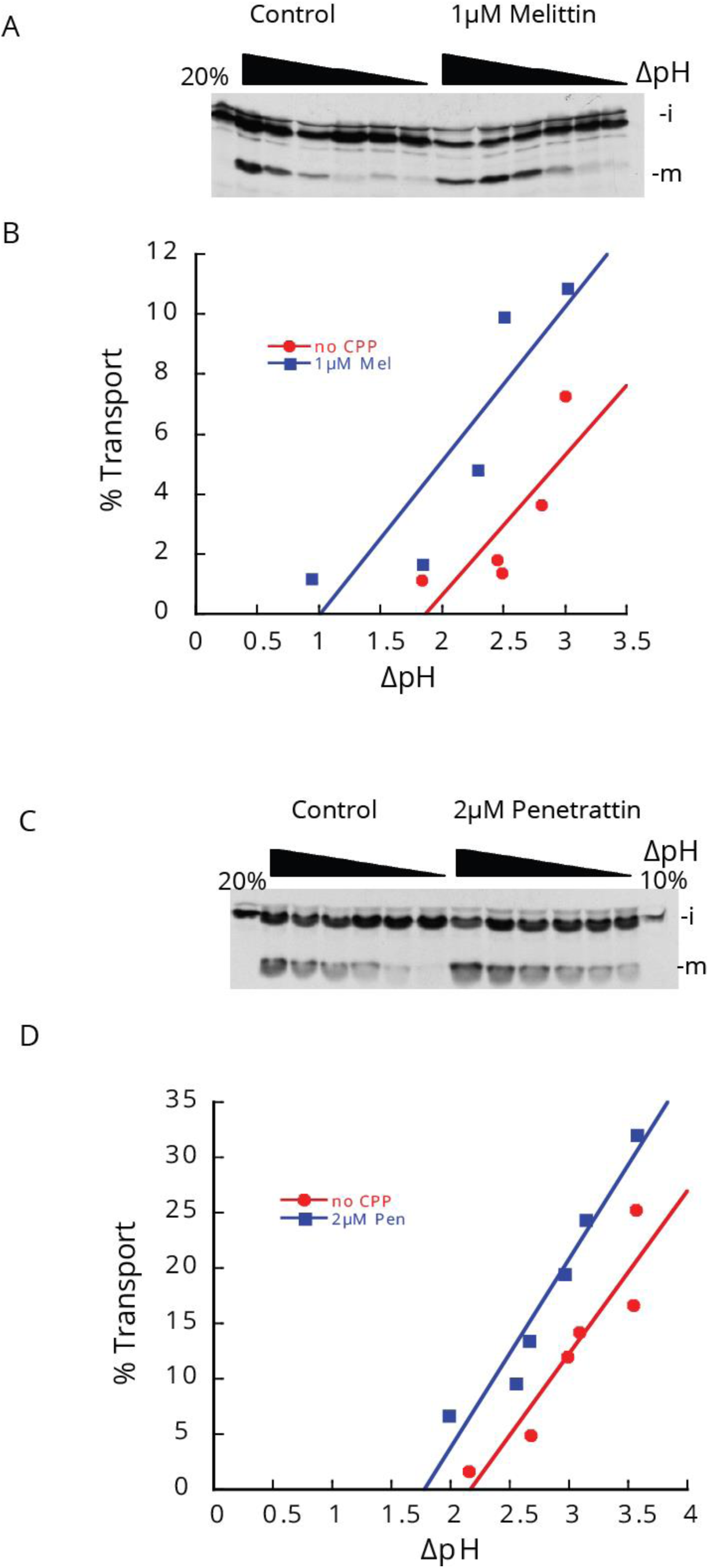
CPPs lower the ΔpH threshold for Tat transport. ΔpH measurements were made by observing the quenching of NED fluorescence by application of actinic light of varying intensities. Transport reactions were stopped with 1mM nigericin and valinomycin immediately followed by centrifugation and resuspension in SB. Protein transport was measured as in Fig. 1.

### CPPs interact with the thylakoid membrane, causing a change in density

The ΔpH and Δψ measurements and the ΔpH threshold experiments suggest that CPPs change the structure of the thylakoids. We suspected that the thylakoids were reorganized in response to interaction with the CPPs. Previously it has been observed that the vesicle inducing protein in plastid1 (VIPP1) stimulates Tat protein translocation and also causes a change in thylakoid density (Lo & Theg, 2012a). We asked whether CPPs could lead to a similar change in the density of the thylakoids. To this end, we performed thylakoid pelleting assays(Lo & Theg, 2012b), observing how much chlorophyll remains in the supernatant after low-speed centrifugation. As seen in Fig. 3, thylakoids generally pellet more readily upon incubation with CPPs, and the response is dose dependent. This suggests that CPPs interact directly with the thylakoid membrane, causing a reorganization of the thylakoid super-structure, which leads to their increased density.

**Figure 3.**
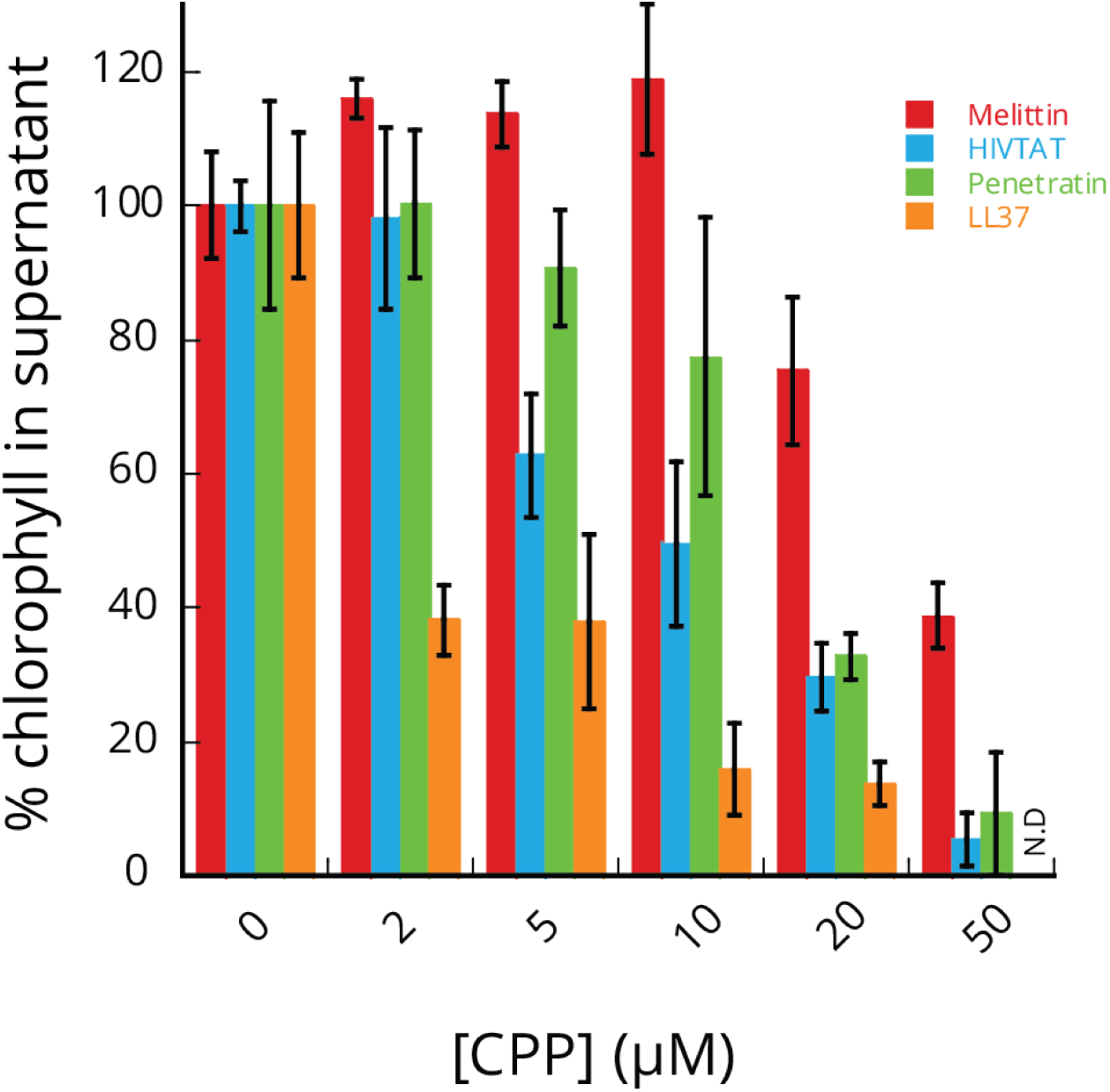
Thylakoids pellet more readily in the presence of CPPs. Thylakoids were incubated with CPPs at indicated concentrations in the dark for 10 min. Thylakoids were then centrifuged at 100 rcf for 1 min. The supernatant was then diluted tenfold into 89% Acetone. Chlorophyll concentration was determined spectroscopically.

### CPPs stimulate Tat transport in bacterial inverted membrane vesicles

It is widely held that the mechanism of protein transport on the Tat pathway is the same in all Tat systems found in the different domains of life (Celedon & Cline, 2012a, Palmer & Berks, 2012), and this view has gained further support from recent work in our lab (Zhou et al., 2023). Accordingly, we would predict that the stimulation of the Tat pathway by CPPs demonstrated above in chloroplasts would apply to the bacterial Tat pathway as well. To test this, we examined the effect of CPPs on the transport of the model substrate pre-SufI-IAC into inverted membrane vesicles (IMVs) prepared from JM109 cells overexpressing TatABC(Bageshwar & Musser, 2007, Bageshwar et al., 2009). As in thylakoids, we again found that CPPs stimulated Tat transport in the bacterial system (Fig. 4). This result suggests that CPPs can act universally in stimulating Tat transport, implying a shared mechanism across systems. The CPP stimulation effect is lost at higher concentrations of CPPs, likely due to pmf uncoupling as in thylakoids (Fig. 1E,F).

**Figure 4.**
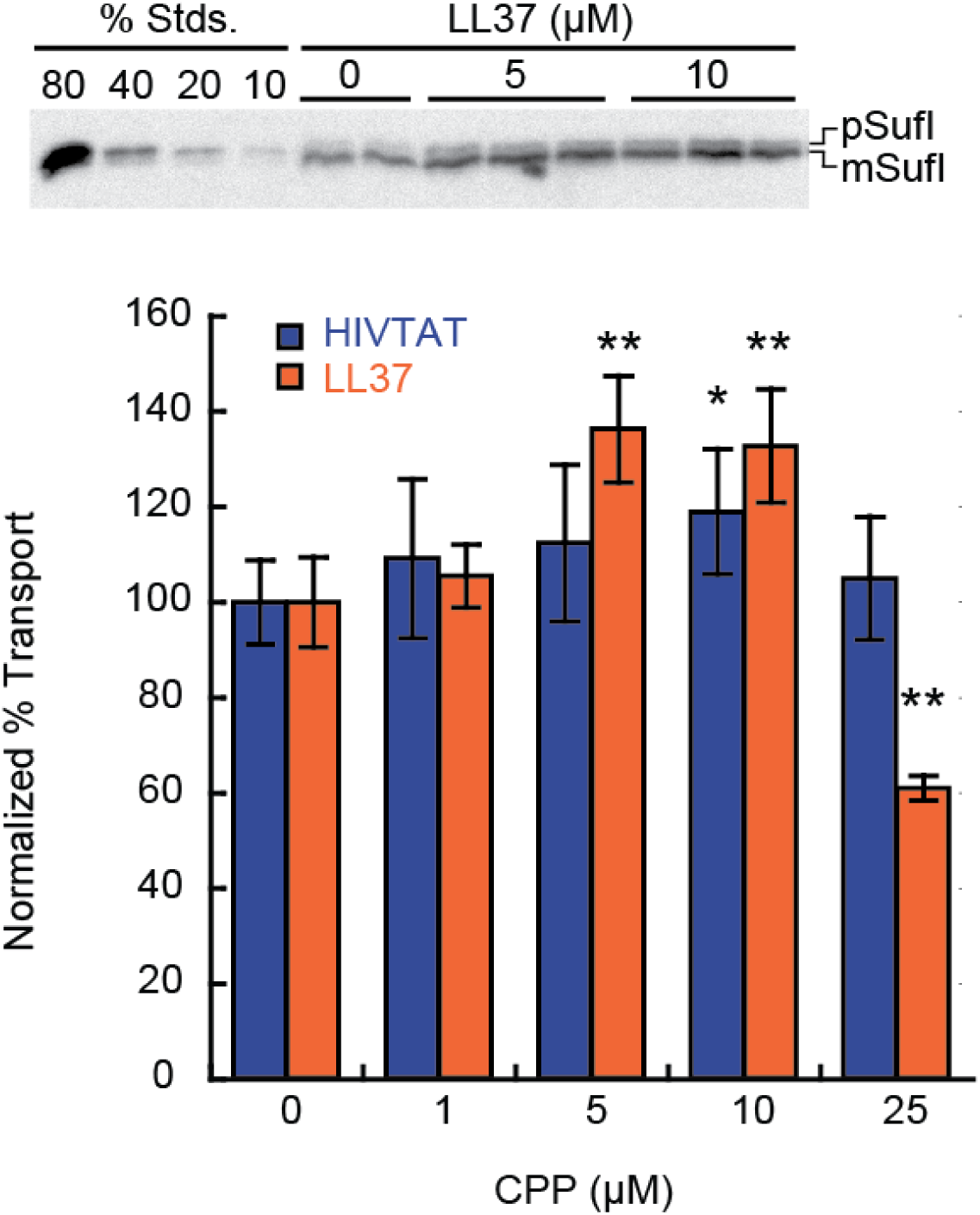
Stimulation of Tat transport in E. coli IMVs by CPPs. Biotinylated pre-SufI-IAC substrate protein was transported into IMVs at 37 °C with indicated concentrations of CPPs. Reactions were stopped after 8 min on ice. Samples were treated with Proteinase K and mature protein was analyzed by SDS-PAGE and α-biotin Western blotting. Error bars indicate SD (n≥4). Significance of CPP effect was determined by ANOVA post-hoc analysis (*p<0.05, **p<0.01).

### The amphipathic domains of Hcf106 stimulates Tat protein translocation

One of the biophysical characteristics linking different CPPs is their abilities to form amphipathic helices. All TatA- and TatB-family member proteins, including Tha4 and Hcf106, also have amphipathic helices, essential for their function, and the APH of Hcf106 is particularly enriched in arginines, as are many CPPs. Given the similarities of these APHs to CPPs we tested their ability to stimulate translocation. While the Tha4 APH peptide did not increase transport of iOE17 (Fig. 5A), the Hcf106 APH peptide did stimulate translocation at a level comparable to CPPs (Fig. 5E). As with the CCPs, we noted a decline in transport stimulation at 20 µM Hcf106 APH peptide. However, unlike the situation with CCPs, we observed that the Hcf106 APH peptide neither compromised membrane integrity nor increased the thylakoid membrane density at this concentration (Fig. 5B-D). The reason for this decrease in iOE17 transport at 20 μM Hcf106 APH peptide remains unknown; we did not probe this phenomenon further.

**Figure 5.**
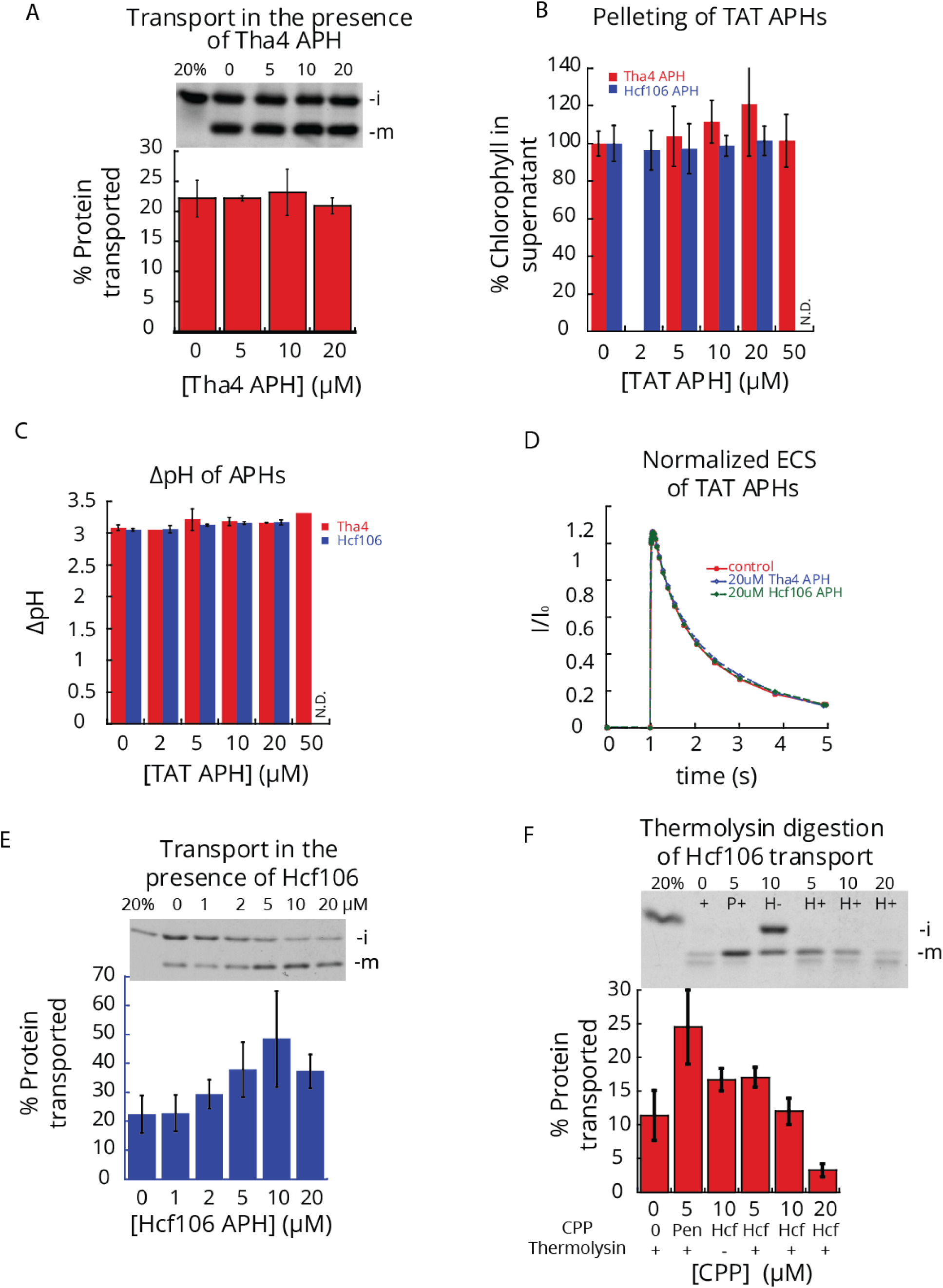
Transport of iOE17 in the presence of Tat APHs. A) As in Fig. 1, thylakoids were incubated with the Tha4 APH peptide and transport was performed. B) Pelleting assays were performed as in Fig. 3. C) ΔpH analysis was performed as in Fig. 1. D) ECS measurements were performed as in Fig. 1. E) As in Fig. 1, thylakoids were incubated with Hcf106 APH and transport was performed. F) Transport reactions were thermolysin treated as indicated (+). Protease was quenched with 25mM EDTA. Thylakoid reisolation and SDS-PAGE followed as in Fig. 1.

Interestingly, the mature substrate protein formed in the presence of the Hcf106 APH peptide at 10 and 20 µM was not fully protected from digestion with thermolysin, suggesting that some fraction had not completely crossed the thylakoid membrane (Fig. 5F). The surprising appearance of a matured but protease-sensitive Tat substrate protein is not new in the literature but is very unusual(Di Cola & Robinson, 2005, Frobel et al., 2012, Leheny et al., 1998). Fröbel *et al*.(Frobel et al., 2012) proposed that in the bacterial system a precursor could be bound by TatC in such a manner that allows the signal peptide to cross the membrane without full translocation of the passenger protein. Whether we are observing the same phenomenon here and what role the Hcf106 APH peptide plays in it, remain to be elucidated.

### Membrane thickening abolishes Tat transport

Having established that CPPs and the Hcf106 APH peptide stimulate Tat protein transport, presumably through membrane thinning and destabilization, we next determined the effect of membrane thickening. To thicken, or in other words strengthen, the thylakoid membrane, we employed trifluoroethanol (TFE). TFE inserts into the membrane at the hydrophobic/hydrophilic interface and increases the lateral membrane pressure, which in effect stabilizes the lipid bilayer(van den Brink-van der Laan et al., 2004, Yuan et al., 2017). We found that Tat transport is very sensitive to TFE, with a precipitous drop in transport at 1% (v/v) TFE (Fig. 6). The ΔpH was not greatly affected by TFE, ruling out the possibility of an indirect effect of TFE on transport through a loss of the pmf. As before, to compare an effect on Tat transport to that with a proteinaceous pore translocon, we examined the effect of TFE on Sec transport. TFE did reduce Sec transport, but still supported Sec transport at concentrations 2.7-fold higher than that required to abolish Tat transport (Fig. 6). Considering the complementary effects of TFE and CPPs, along with reports on lengthening the TatA and TatB TMHs (Hao et al., 2022a, Mehner-Breitfeld et al., 2022, Stockwald et al., 2022), it is apparent that membrane structure, in particular, membrane thinning, is critical for the Tat translocation mechanism.

**Figure 6.**
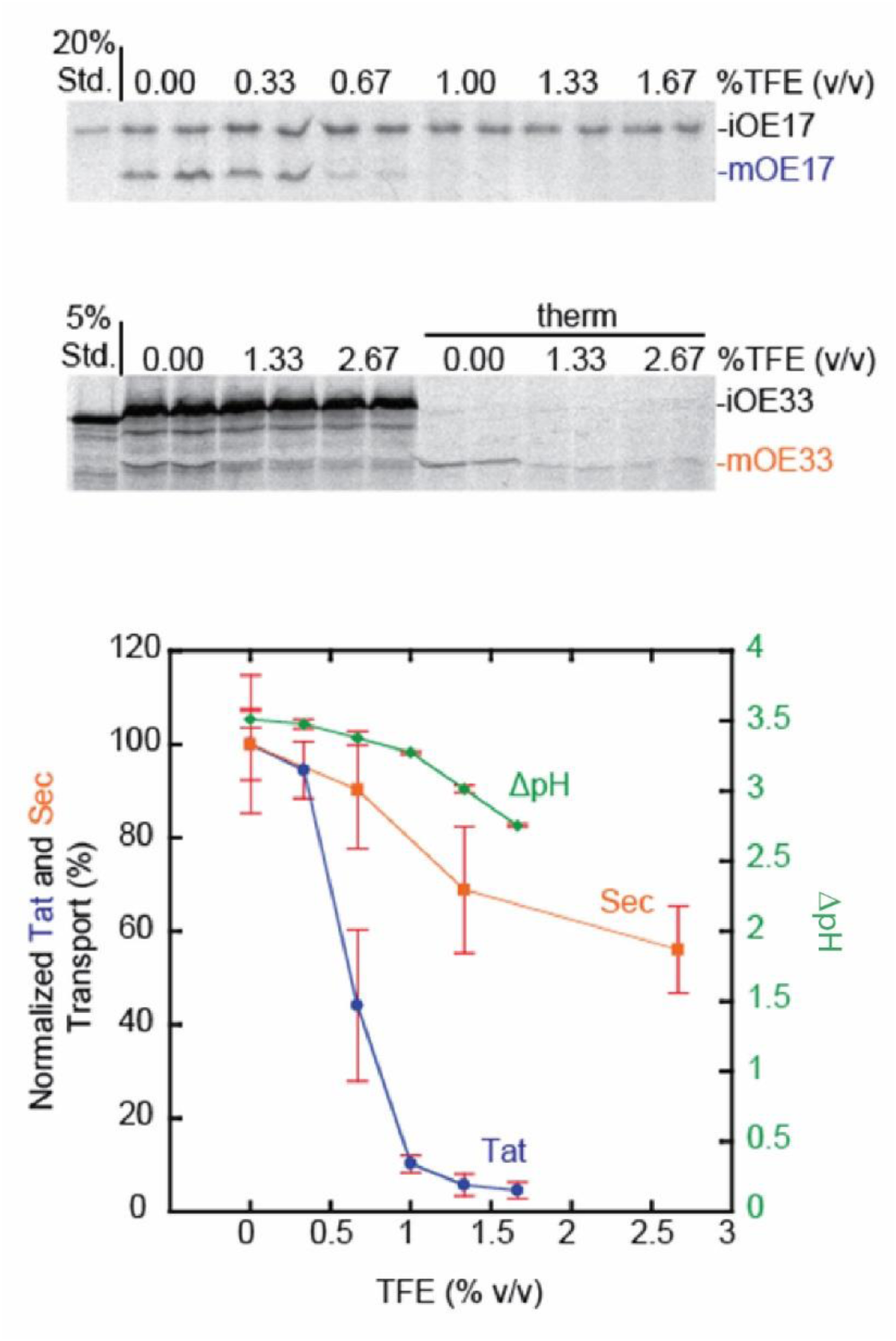
Inhibition of Tat transport by TFE. Thylakoids were assayed for Tat (iOE17, blue) or Sec (iOE33, orange) transport in the presence of indicated TFE concentrations. Thylakoids were supplemented with ethanol such that the sum total concentration (%v/v) of TFE and ethanol was equivalent in all reactions. Post-transport samples were treated with thermolysin (therm) where indicated and analyzed by SDS PAGE and fluorography. Mature bands were quantified (n≥3, SD). Thylakoid ΔpH was measured at indicated TFE concentrations (green, n=3, SD) as described previously by 9AA fluorescence quenching (Theg & Tom, 2011).

## Discussion

In this work, we demonstrated that CPPs stimulate Tat transport (Fig. 1) by interacting with and changing the thylakoid membrane (Fig. 3) and thereby lowering the ΔpH threshold for translocation (Fig. 2).

Since CPPs are known to cause thinning in membranes(Chen et al., 2003, Herce & Garcia, 2007, Madani et al., 2011, Mecke et al., 2005), it is likely that they also act to stimulate the Tat pathway by thinning the thylakoid membrane. This mechanism also successfully predicts the decrease in the threshold ΔpH for protein transport mediated by melittin and penetratin. Given that the bacterial Tat system is also stimulated by CPPs, we can extrapolate that the membrane thinning mechanism of action is conserved across systems. This concurs with recent observations that *E. coli* TatA causes membrane defects in coordination with the substrate protein(Hou et al., 2018).

Membrane thinning provides an interesting mechanism for translocation in the context of coupling of the pmf to protein transport. Assuming a random distribution of CPPs in the thylakoid membrane, the global thinning caused by the pmf(Kirchhoff et al., 2011) is likely mimicked by the action of the CPPs. This idea is further supported by CPP stimulation of the bacterial pathway because CPP interaction with model bacterial membranes is more widely studied(Lee et al., 2005). We would expect that the thinning could reach an extreme locally as the translocon subunits each contribute to narrowing the membrane through hydrophobic mismatch in their TMD regions (Hao et al., 2022a, Mehner-Breitfeld et al., 2022, Stockwald et al., 2022) and/or potential contributions from their APH regions. Should this local thinning draw further Tha4 subunits into the vicinity, then the membrane destabilization calculated by Rodrigiez et al.(Rodriguez et al., 2013b) that occurred when the TatA mulitmer exceeds nine subunits would provide the conditions for the formation of a toroidal pore. However, one could expect the Tat machinery to induce controlled pores that would not result in massive ion leakage(Asher & Theg, 2021, Teter & Theg, 1998) by forming a sphincter, having a very short life, or some combination of these two.

In the toroidal pore model, the head groups of lipids would be expected to interact with the substrate. While lipid-substrate interactions have been demonstrated(Bageshwar et al., 2009, Rodriguez et al., 2013b), cross links between the two have not been investigated and may be a worthy subject of study to determine which lipids comprise the toroidal pore during transport, since some lipids are essential to Tat transport (Rathmann et al., 2017). One would expect to find the pore consisting of lipids that introduce curvature in the membrane. Interestingly, branched and phospholipids of the thylakoid membrane tend to interact with photosynthetic membrane proteins (Mizusawa & Wada, 2012). If this data can be extrapolated to more membrane proteins, then TatA recruitment may serve as a means of collecting the lipids that induce the curvature necessary for toroidal pore formation.

A further driver for membrane thinning could be provided by attraction of the charged arginine residues in the Tat substrates’ signal peptide to the trans-side of the membrane, as is hypothesized for the mechanism of membrane crossing for arginine-rich CPPs (Herce & Garcia, 2007). However, this view is tempered by the recognition that the signal sequence arginines are likely buried in the receptor complex and interacting with TatC (Rollauer et al., 2012, Zoufaly et al., 2012). Thus, if they play a role, it may be upon release of the precursor from the receptor complex. Substrate proteins poised at the pore site could then traverse the membrane while the pore is open with the signal acting as an anchor. Such pores would be expected to close quickly once the substrate is cleared as the membrane-thinning pmf would be locally decreased due to the concurrent movement of protons through the pore. This would be a natural consequence of the large number of protons observed to be released from the proton gradient per protein translocated(Alder & Theg, 2003).

A toroidal pore itself does not provide directionality. Since transport occurs *in vitro* without soluble factors, directionality would be provided by the translocon. Alternatively, substrate exit from the translocon to the trans side of the membrane could be random. In the case of a failed translocation attempt, rebinding of the substrate and reassembly of the active translocon may be necessary. We recognize that a mechanism requiring multiple attempts might explain, in part, the extraordinarily high energetic cost(Alder & Theg, 2003) and time(Celedon & Cline, 2012b, Musser & Theg, 2000, Teter & Theg, 1998) of the translocation reaction, as only successful events would be scored as protein transport..

Here, we have not directly addressed whether the Tat pore is proteinaceous or toroidal. However, membrane thinning is fundamental to toroidal pore formation, which is generally not the case for proteinaceous pores. An example of an exception would be proteinaceous pores formed by mechanosensitive ion channels(Louhivuori et al., 2010). Therefore, it remains possible that membrane thinning causes a conformational switch in TatA that induces assembly of a proteinaceous pore. Again, we disfavor this model due to the problem of the substrate contacting the hydrophobic TMHs of the Tat machinery. A solution to the substrate interaction with TMHs has been proposed in the charged zipper model wherein the C-terminal domain of the TatA family proteins are proposed to line a TatA pore(Walther et al., 2013). This model is not supported by the evidence {Alcock, 2017 #26358;Rodriguez, 2013 #26297;Berks, 2015 #26291}.

It is interesting that the Hcf106 APH peptide caused Tat transport stimulation, while the Tha4 APH peptide did not. This may be because Tha4 APH may require a higher concentration to act on the membrane. Indeed, an actively transporting Tat translocase contains a higher stoichiometry of Tha4 than Hcf106(Celedon & Cline, 2012b) and only one Hcf106 subunit within the complex may be involved per substrate transported. Additionally, the full length Tha4 APH may be more active than the shorter peptide we used, despite having a reduced hydrophobic moment. We also note that the APH of That4 has one arginine, whereas the APH of Hcf106 has five. This residue is found in abundance in CPPs, which have in fact been referred to as arginine-rich peptides (Futaki et al., 2013, Hao et al., 2022b); a titration showed that the optimal number for maximum CPP activity is eight arginines (Futaki et al., 2001).

In summary, we have shown here that protein translocation on the Tat pathway is stimulated by CPPs and CPP-like peptides (the Hcf106 APH) in both thylakoids and bacteria. We interpret this as additional evidence that Tat protein transport involves membrane thinning and ultimately lipid bilayer breakdown to form toroidal pores which allow protein substrates to traverse the biological membranes. The transient and local nature of such pores has rendered them undetectable to date, and this will be the challenge for our future studies of this enigmatic pathway.

## Materials and Methods

### Reagents

Cell Penetrating Peptides were purchased from Anaspec. Tha4 APH (KKLPEVGRSIGQTVKSFQQAAK) and Hcf106 APH

(KGLAEVARNLGKTLREFQPTIREIQDVSREFKSTLER) were purchased from Lifetien at >90% purity. Peas were purchased form Harris Seed.

*Buffers.* Grinding buffer (GB): .05M (K)HEPES, .33M Sorbitol, 1mM MgCl_2_, MnCl_2_, 2mM Na_2_EDTA, pH to 7.3 then .1%BSA. Import Buffer (IB): .05M K-Tricine, .33M Sorbitol, 3mM MgCl_2_ pH 8. 2x Sample Buffer (2xSB): .125M Tris (pH 6.8), 4% SDS, 20% glycerol, .005% Bromophenol Blue, 10% BME.

Methyl viologen (MV), valinomycin, N,N Naphthylethylene diamine (NED), Phosphatidyl glycerol 18:1-06:0 2- (4-nitro-2,1,3-benzoxadiazol-7-yl) (PG-NBD), and nigericin were stored in 100% EtOH.

### Plant growth and thylakoid isolation

Peas were grown on soil as described previously (Lo & Theg, 2011). Pea leaves and shoots from 10 to 14-day old plants were blended in short pulses in GB. Blendate was centrifuged for 5 min at 4250 rpm. The pellet was resuspended in GB and passed through a Percoll gradient (50:50 percoll:2xGB). The lower band of intact chloroplast was then washed twice with IB. Thylakoids were obtained by lysing chloroplasts in water supplemented with 5 mM MgCl_2_. Lysis was stopped by adding an equal volume of 2xIB. Isolations were quantified by chlorophyll content as described(Lo & Theg, 2011).

### Thylakoid Tat Translocation assays

*Translation.* In vitro translations were performed with wheat germ extract as directed by the manufacturer (Promega or tRNA Probes) with ^3^H leucine or ^35^S methionine. Plasmid encoding iOE17 was described previously(Alder & Theg, 2003).

*Translocation.* Transport reactions (60 µl) contained 2 µl translation product, CPPs at indicated concentrations, and 0.33 mg/mL chlorophyll-equivalent thylakoids in 1xIB. Reactions were initiated by addition of thylakoids, incubated 6 min at room temperature under illumination, and stopped with 10 volumes of cold IB and transferred to the dark on ice. Thylakoids were reisolated by pelleting at 3,000 rpm for 5 min. The pellet was resuspended in SB, boiled for 10 min, and analyzed by SDS-PAGE and fluorography.

*Thermolysin digestion.* After stopping a transport reaction, 200 μg/mL thermolysin and 5 mM CaCl_2_ were added where noted. After a 30-minute incubation on ice, EDTA was added to a final concentration of 25 mM. Reisolation of thylakoids and proceeding followed as above.

### Thylakoid Sec Translocation assays

A plasmid containing the thylakoid Sec substrate iOE33 was a gift from Ken Cline; the protein was in vitro translated as described above. SecA was purified from plasmid pET28a-cpSecA1 as described(Endow et al., 2015).

Thylakoids were pretreated with SecA for 15 min at room temperature. Transport reactions (60 µl) containing 4 µl radiolabeled substrate, 5 mM ATP, 25 µg/mL SecA, and 0.33 mg/mL thylakoids were illuminated at room temperature for 30 min. Samples were subsequently treated and analyzed as described for Tat transport.

#### Measurement of Δψ

Measurements of the Δψ-reporting carotenoid electrochromic shift (ECS) were performed on a JTS-10 spectrophotometer. MV (20 μM), thylakoids (20 µg Chl), the given concentration of melittin were brought to a total volume of 1mL in IB. The rate of decay of the ECS signal was monitored after delivering two 9 ms flashes 100 ms apart.

#### Measurement of the thermodynamic threshold for Tat protein transport

ΔpH measurements were made on a Horiba Flouromax fluorometer. 20 mM MV, 4 mM NED, 10 μL of radiolabeled precursor, 120 µg thylakoids, and the given concentration of CPPs were made up to a 2 mL total volume with IB in a 2 mL microcentrifuge tube. After 10 s of dark the excitation beam was applied. After 1 minute from experiment start, an actinic light of varying intensity, depending on desired ΔpH, was applied for 6 minutes. The transport reaction was stopped with 1 mM final concentrations of nigericin and valinomycin. Thylakoids were immediately reisolated, resuspended in SB and analyzed by SDS-PAGE and fluorography.

#### Measurement of relative thylakoid density

As in Lo and Theg(Lo & Theg, 2012b), 60 µL of 0.33 mg/mL chlorophyll-equivalent thylakoids were incubated with CPPs in the dark at ambient temperatures for 10 min and then centrifuged at 100 rpm for 1 min. 20 µL of supernatant was removed and diluted into 180 µL 89% acetone and the concentration of chlorophyll was calculated from absorbance as previously described(Lo & Theg, 2012b).

### E. coli inverted membrane vesicle (IMV) transport

The pre-SufI-IAC protein was overexpressed in *E. coli* from plasmid pSufI-IAC(Bageshwar et al., 2009), which encodes SufI precursor with substituted native cysteines and a single C-terminal cysteine. Protein was purified under native conditions as previously described for pre-SufI(Bageshwar & Musser, 2007). Eluted protein (6.5 µM) was biotinylated by addition of 0.7 mM TCEP and 500 μM biotin maleimide and incubation on ice for 30 min. The labeling reaction was quenched with 2 mM DTT.

IMV isolation from JM109 overexpressing pTatABC and transport assays were essentially as previously described(Bageshwar & Musser, 2007). Briefly, 110 nM biotinylated pre-SufI-IAC was mixed with IMVs (A_280_=5), 4 mM NADH, and indicated concentrations of CPPs. Reactions were incubated at 37°C for 8 min and stopped on ice. Samples were subsequently treated with 0.3 mg/mL Proteinase K for 40 min at RT, quenched with 3 mM PMSF on ice, and diluted with an equal volume of 2xSB. Samples were analyzed by SDS-PAGE and α-biotin Western blotting as described previously(Ganesan et al., 2018).

## Acknowledgements

We thank Siegfried Musser and Shruthi Hamsanathan for pSufI-IAC, Tracy Palmer and Jon Cherry for JM109 expressing pTatABC, and Ken Cline for the iOE33 clone. We gratefully acknowledge support from the Division of Chemical Sciences, Geosciences, and Biosciences, Office of Basic Energy Sciences of the U.S. Department of Energy through Grants DE-FG02-03ER15405 and DE-SC0020304 to SMT.

## Author Contributions

**Robert McNeilage:** Conceptualization; performed experiments in Figures 1, 2, 3 and 5; writing – original draft, review and editing. **Iniyan Ganesan:** Conceptualization; performed experiments in Figure 3 and 6; writing – original draft, review and editing. Johnathan Keilman: Performed experiment in Figure 6; writing – review and editing. Steven M. Theg: Conceptualization; supervision; funding acquisition; project administration; writing – original draft, review and editing.

## Disclosure and competing interests statement

The authors declare that they have no conflict of interest.

## Data Availability

This study includes no data deposited in external repositories.

## Notes

### Competing Interest Statement

The authors have declared no competing interest.

### Summary of Updates

We fixed numerous grammatical errors and improved readability of the paper

